# Tuning nanoscale lipid clustering controls influenza viral binding

**DOI:** 10.64898/2026.09.28.755053

**Authors:** Steinar Mannsverk, Shreyaa Srinivasan, Diogo Filipe, Frederick A. Heberle, Ana M. Villamil Giraldo, Peter M. Kasson

## Abstract

Influenza virus attaches to cells by binding to sialylated glycans. Subsequent endocytosis of the virus and endosomal maturation leads to a conformational change in the hemagglutinin coat protein that ultimately results in membrane fusion and release of the viral genome into the host cytoplasm, where it is trafficked to the nucleus and commences replication. Individual hemagglutinin-glycan binding affinities are weak (K_d_ approximately mM), so multivalency is key to viral attachment. Receptor density or clustering has previously been shown to affect influenza binding, but the overall spatial heterogeneity of both plasma and endosomal membranes has not been fully addressed. Here, we show that nanoscale lipid clustering can increase local concentration of glycosphingolipids and greatly enhance influenza viral binding in a way that macroscopic phase separation does not, and that *binding avidity is dependent on the spatiotemporal scales of this clustering*. This clustering is driven by sphingolipids and cholesterol and the nanoscale lipid environment of these clusters persists to affect the viral entry process. This finding demonstrates how fine receptor patterning can drive infection and illustrates the importance of tuning spatial scale in liquid-liquid phase separation.

## Introduction

Influenza A virus (IAV) binds to sialic-acid-terminated glycans on the cell surface. These potentially include glycosphingolipids, O-linked glycans, and N-linked glycans. Both glycolipid and glycoprotein candidate receptors can undergo clustering and potentially phase separation, but glycosphingolipids are most often used as model receptors for influenza virus because of their relatively straightforward chemical reconstitution. Individual hemagglutinin-glycan affinities are in the mM range ^1^, yet multivalent effects in viral attachment mean that viruses can stably attach to the cell surface and bind multivalent synthetic substrates with nanomolar EC_50_ values ^2-4^. While receptor clustering has been shown to affect viral binding ^5^, macroscopic phase separation does not appear to be critical for influenza entry itself ^6^. One caveat is the potential role of EGFR in signaling for viral endocytosis ^7, 8^, but this is not a first-order binding effect.

Glycosphingolipids contain a sphingosine moiety and are expected to co-segregate with sphingomyelin. Both longstanding and recent work have demonstrated that sphingomyelin can associate with cholesterol and can also drive liquid-liquid phase coexistence in conjunction with a low-Tm phospholipid, such as POPC ^9-12^. Depending on the mixture composition and the temperature, the length scales for phase coexistence can either be macroscopic (micron or larger) or nanoscale (as small as 2-7 nm). Sphingomyelin exists at extremely high mole fractions in the outer leaflet of mammalian plasma membrane—29 mol % in human red blood cell outer leaflets^13^—but its abundance decreases dramatically through the endosomal pathway, reaching∼10-15 mol % in mid-late endosomes where influenza membrane fusion occurs.

In lipid membrane mixtures that undergo liquid-liquid phase separation, there is not an abrupt transition from an *ideal* single-phase mixture to two coexisting liquid phases as a function of lipid composition. Instead, the chemical interactions that drive liquid-liquid phase separation result in non-ideal mixing within a macroscopically single-phase mixture. Compositional fluctuations are predicted to increase in spatial and temporal scale until the phase boundary is reached ^14, 15^. The physical interactions driving this behavior are the same as those responsible for phase separation, just operating to drive inhomogeneities within the macroscopically single-phase region.

Here, we use single-virus binding and fusion assays to measure the effect of lipid composition on influenza viral entry. This family of assays uses fluorescent probes to track viral movement, binding, and both lipid and content mixing in the fusion process. Variants of this approach have been used to measure viral entry within cells as well as binding and fusion to bilayers, plasma membrane vesicles, liposomes, and isolated endosomes. Here, we opt for synthetic membrane partners to maximize the chemical controllability of the system and titrate membrane components as precisely as possible.

Using this approach, we show that influenza binding to glycosphingolipid receptors has a non-monotonic dependence on sphingomyelin in the target membrane. Binding avidity peaks at approximately 10 mol % sphingomyelin in the presence of 20 mol % cholesterol and 2 mol % GD1a receptor. This is dependent on both cholesterol and the sphingolipid moiety of the receptor - if either is removed, then no increased binding is observed. This appears to coincide with a nanometer-scale length regime for clustering of sphingolipids and cholesterol well below the phase boundary for macroscopic liquid-liquid phase coexistence. The nanoscale organization of lipid composition that drives viral binding then persists to affect the fusion process. This suggests that viral binding may be influenced by and in turn stabilize a local lipidome that determines the next stage of viral entry and infection.

## Results and Discussion

### Influenza binding avidity is highly responsive to sphingomyelin abundance

Prior work has shown that influenza binding avidity is increased by the addition of cholesterol to a phospholipid-glycosphingolipid mixture and that binding also has a cooperative dependence on glycosphingolipid concentration with a Hill coefficient of ∼2.5 ^5^. We tested the dependence of influenza binding to the same lipid mixture but with the addition of 5-20 mol % egg sphingomyelin (predominantly d18:1/16:0 and abbreviated as SM). In this case, we used a similar microfluidic binding assay but with target liposomes immobilized rather than a target bilayer, and influenza virions were quantified using immunofluorescent labeling of hemagglutinin (HA) glycoproteins (Fig. 1a). Overall, 81±3% of HA^+^ particles colocalized with a fluorescently labeled liposome, indicating that the majority of viral particles were specifically bound to a liposome. We observed a notable increase in influenza binding avidity at 10 mol% SM (Fig. 1b-c). Surprisingly, avidity decreased again at ≥20 mol % SM (Fig. 1c). Because the closest ternary mixture analogue, POPC : Chol : SM, shows nanoscale liquid-liquid phase coexistence at these compositions, we suspected an effect related to liquid-liquid phase separation. Our optical imaging of bilayers (Fig. S1) and prior imaging studies on giant unilamellar vesicles ^11^ both do not show detectable liquid-liquid phase separation at 10 mol % SM and 37 ºC, nor do SANS studies. However, FRET does indicate phase coexistence in similar mixtures ^16, 17^. By comparing the spatial sensitivities of these techniques, it has been concluded that the length scales for phase separation are likely ∼10 nm. Since GD1a glycosphingolipids share a sphingosine moiety, they would be expected to partition into Lo phases with sphingomyelin. Such an avidity-enhancing effect of nanoscale clustering is compatible with other GD1a clustering-induced effects on viral binding driven by other physical mechanisms. Consistent with this, we did not observe increased binding avidity when GD1a was substituted with a DNA-lipid attachment factor ^18^ that partitions into liquid disordered (Ld) phases (Fig. 1c), again suggesting an enhancement related to liquid-ordered phenomena.

**Figure 1.**
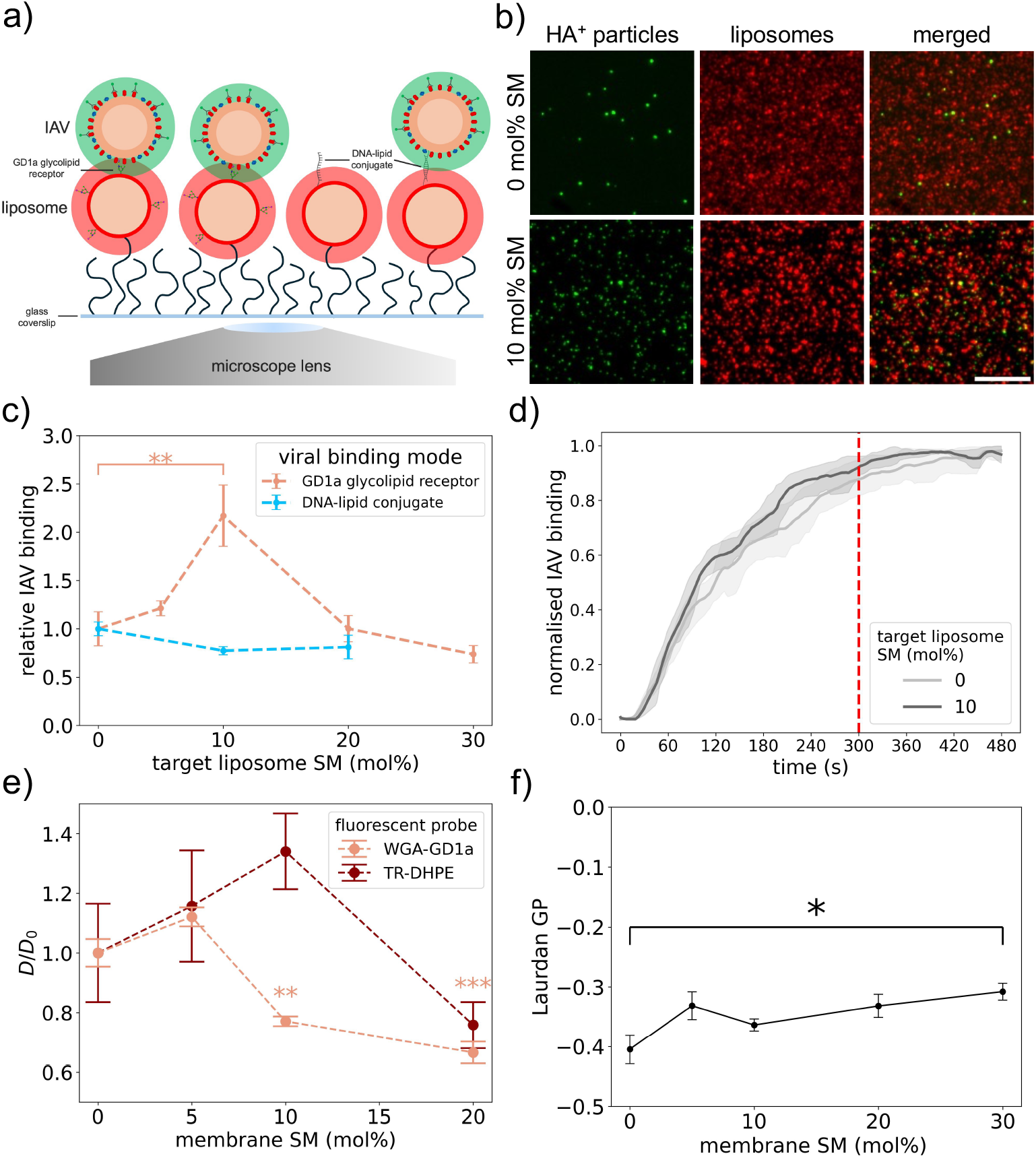
Influenza binding to liposomes containing sphingomyelin. **a**) Schematic of the binding assay. Virus is incubated in a flow cell with previously immobilized fluorescently labeled liposomes for 5 min at 37 °C. Unbound virions were washed away, and bound virions were quantified via anti-HA immunofluorescence. **b**) Representative fluorescence micrographs of anti-HA (green) and liposome (red) labeling. The mol % SM is varied between top panels (0 mol %) and bottom panels (10 mol %). Scale bar: 10 µm. **c**) Relative IAV binding to liposomes with the indicated amount of SM, measured as the relative fraction of liposomes with a colocalized HA-labelled virion. **d**) Binding time course showing influenza binding after viral flow-in relative to maximal binding. Red vertical dashed line indicates the time when unbound virions were removed in panes **b-c**. Shaded regions represent 90% bootstrapped confidence intervals. No difference in time course was detected between SM compositions using a bootstrapped permutation Kolmogorov–Smirnov test performed on individual channels (n = 1000, p = 0.21). **e**) Change in lipid diffusion coefficients for GD1a labelled with WGA-iFluor 594 or the lipid probe TR-DHPE, estimated from FRAP measurements on supported lipid bilayers. Data are plotted as D/D_0_, where D_0_ denotes the diffusion coefficient for 0 mol % SM. Error bars show standard error. **f**) Laurdan generalized polarization (GP) values as a function of mol % SM, where higher GP indicates lower membrane fluidity. All liposomes contained 20 mol % DOPE, 20 mol % chol, 1 mol % DPPE-Biotin, 2 mol % GD1a or DNA-lipid conjugate (where stated), and POPC to 100 mol %. In addition, liposomes in panels **b**-**d** contained 0.5 mol % DOPE-Atto 647. Statistical significance in **c, e**, and **f** was assessed using one-way ANOVA followed by Tukey’s HSD post hoc test (* *p* < 0.05, ** *p* < 0.01, *** *p* < 0.001).

### Avidity-enhancing membranes remain fluid and single-phase on the micron scale

To probe this avidity-enhancing effect, we first verified that the single-point measurements of binding do not bias avidity estimates. Influenza virus reached >90% saturation in binding to 0 and 10 mol % SM after 300 s, the time interval used for avidity estimates (Fig. 1d). Similarly, micron-scale diffusion of GD1a showed a substantial slowing from 0-5 mol % SM (unchanged) to 10-20 mol % SM content (20-30% slower, Fig. 1e). In the same mixtures, no significant change was seen in the diffusion of Texas Red-DHPE (Fig. 1e). The baseline diffusion coefficient estimates from FRAP of these two probes were 2.1 µm^2^/s and 0.7 µm^2^/s respectively; the slower estimated diffusion of TR-DHPE is likely due to the fact that this dye is incorporated into both leaflets of the glass-supported bilayer whereas the WGA only binds to the solute-facing leaflet. Prior studies have shown that diffusion is slower in the support-proximal leaflet, although estimates vary widely from 17% to 50-fold ^19, 20^.

These results allow us to exclude two hypotheses for lipid organization in these membranes. First, the micron-scale diffusion observed is in the range for single-phase liquid bilayers or Ld phase rather than solid phase. Second, the change in diffusion coefficient observed over this range of compositions limits the hydrodynamic radius of the diffusing GD1a via the Stokes-Einstein relation. Since the diffusion coefficient changes by less than 50%, GD1a must diffuse on the length scale of the bleaching spot with a hydrodynamic radius similar to GD1a in the absence of SM. This implies that the diffusing species is not a liquid-ordered (Lo) phase or large complex. We thus infer that if there are Lo regions that GD1a partitions into, it undergoes relatively fast exchange between such ordered regions and the surrounding lipid environment (Ld or otherwise) on the length time scales of photobleaching recovery, thus accounting for the measured diffusion coefficients. Laurdan GP values also showed a slight but steady increase in membrane ordering from 0 to 30 mol % SM (Fig. 1f). All these measurements indicate that 10 mol % SM does not induce a discontinuity in the macroscopic physical parameters of the target membrane.

To further understand this effect of target membrane organization on influenza viral binding avidity, we varied the chemical identity of the components. We tested the omission of sphingomyelin and/or cholesterol as well as the chemical substitution of 5-cholesten-3-one for cholesterol or 18:0 ceramide (Cer) for sphingomyelin. Enhancement of binding was greatest with sphingomyelin and cholesterol (Fig. 2a); cholesterol only had a smaller enhancement consistent with previous reports ^5^. 5-cholesten-3-one did not significantly enhance binding but had a high variance, likely consistent with reports of the formation of gel domains in mixtures of approximately this composition ^21, 22^. Substitution of ceramide for sphingomyelin resulted in an intermediate binding avidity. The phase behavior of this mixture (47 POPC : 20 DOPE : 20 Chol : 10 Cer : 2 GD1a : 1 DPPE-biotin) is likely complex, but the closest ternary mixture analogue shows liquid-gel coexistence at 24 ºC ^22^. Generalization to the more complex mixture used and 37 ºC is non-trivial. Taken together, these results suggest that sphingolipid-cholesterol chemical interactions are necessary for optimal binding. In conjunction with the prior controls, we further hypothesize that maximal binding avidity is achieved when GD1a receptors co-segregate into regions of the membrane with cholesterol and sphingomyelin. These likely result from nanometer-scale lipid clustering.

**Figure 2.**
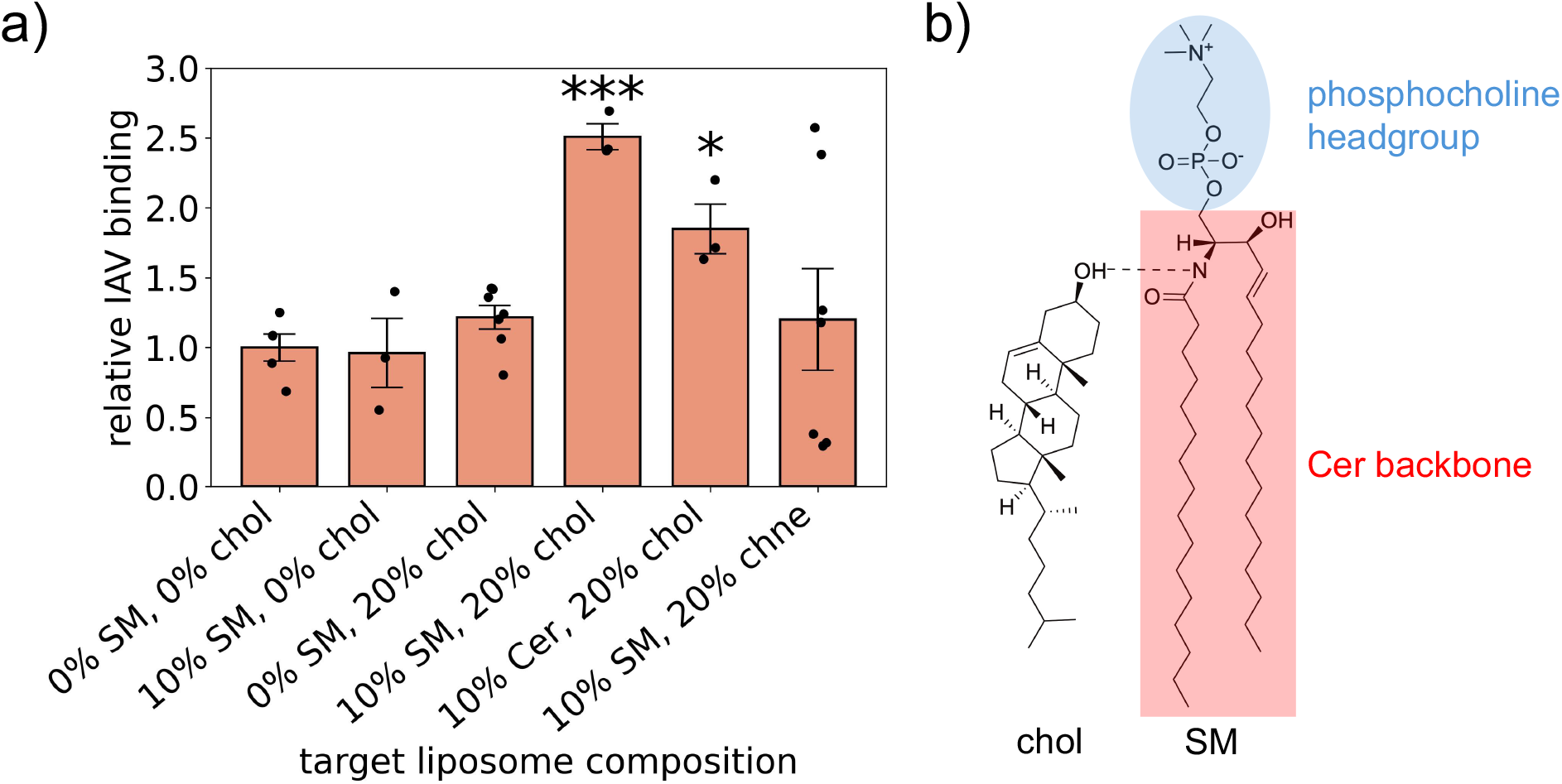
Influenza binding to liposomes with different sphingolipid and sterol components. **a**) Relative IAV binding to liposomes at 37°C as in **Fig. 1a-c**, but with the sphingolipid and sterol components varied. Binding significantly different from the reference condition (0 % SM 0 % chol) is indicated with asterisks (*** p < 0.001, * p < 0.05), determined by pairwise Student’s two-sample t-tests with Bonferroni multiple-hypothesis correction. **b**) Chemical structures of chol (left) and SM (right), highlighting a potential hydrogen bond (dashed line) between the 3β-hydroxyl group of chol and the amide backbone of SM. The SM headgroup and Cer backbone are shown in blue and red, respectively. In contrast, the oxidized chol analogue 3-cholesten-3-one (chne) contains a 3-keto group at the C3-position, which removes the hydroxyl hydrogen-bond donor and is expected to alter, potentially weaken, sterol–SM interactions.

### Simulations show nanoscale clustering increasing with sphingolipid content

To further explore this hypothesis, we conducted molecular dynamics simulations of bilayers containing the key lipid components above and explored their phase and GD1a-clustering behavior. Because the simulations contained ∼1400 lipids each and covered 1-2 µs timescales per replica, they likely do not sample the full set of collective spatial arrangements expected in macroscopic membranes. However, the simulations do sample local lipid-lipid interactions that underlie nanoscale phase separation. To measure these, each simulated composition was analyzed using a Hidden Markov Model (HMM) to classify areas as Lo-like or Ld-like (see Methods). The stability of Lo-like regions increased monotonically from 5% SM : 20% Chol to 10% SM : 20% Chol to 20% SM : 20% Chol (Fig. 3b), supporting the hypothesis of increasing nanoscale cluster stability across these conditions. Similarly, the probability of finding sphingolipid clusters of a larger size increased from 5 to 10 to 20 mol % SM (Fig. 3c) even after controlling for random effects as a function of increasing sphingolipid concentration (see Methods). Both findings are consistent with the expectation that nanoscale clusters should increase in both temporal and spatial scales as the composition approaches a phase coexistence boundary.

**Figure 3.**
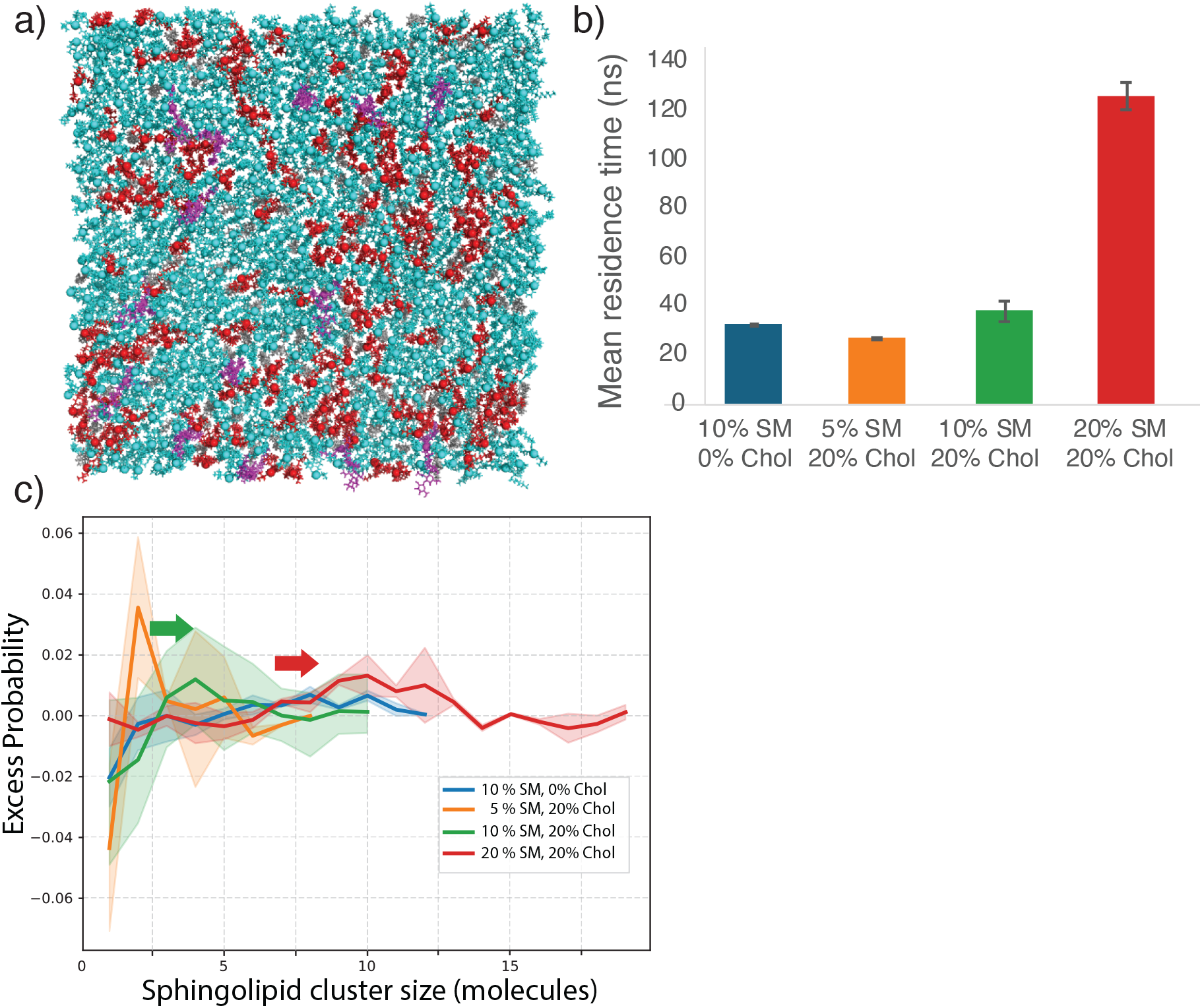
Molecular dynamics simulations of lipid nanoclustering in mixtures replicating experimental conditions. All-atom molecular dynamics simulations were run of POPC:DOPE:SM:Chol:GD1a mixtures replicating the target membranes used for influenza binding experiments. **a**) Representative snapshot showing one leaflet of the bilayer. POPC and DOPE are colored cyan, SM is colored red, GD1a is colored magenta, and Chol is colored off-white. Water and ions are not shown. **b**) Metastability of Lo-like lipids as a function of overall membrane composition. Plotted are decorrelation times for HMM-based classification of lipids as Lo-like or Ld-like (see Methods). Metastability increases with increasing mol % SM. **c**) Excess probability of sphingolipid clusters in simulations of the indicated lipid mixtures. Size distributions of sphingolipid clusters were calculated (see Methods), and excess probability was determined by subtracting the average of 10 random permutations of residue identity. Excess probability indicates increased clustering, and the cluster sizes thus determined increase from 5 to 10 to 20 mol % SM.

Although prior experimental evidence supports favorable thermodynamics for GD1a-GD1a association ^23^, simulated GD1a showed no significant deviation from a random subsample of sphingolipids (p > 0.5, see Methods). If we therefore assume that GD1a positions can be approximated as a random subsample of sphingolipid positions, the simulation results suggest that nanoscale clusters at 10 mol % SM are substantially smaller than at 20 mol % SM. Prior experimental work found that optimal influenza binding of bivalent sialic acid substrates occurred at a ligand-ligand spacing of 55 Å ^24^. Full sampling of patterning on this spatial scale is limited by finite-size effects in our atomistic simulations of bilayer patches approximately 186 x 186 Å, but the trends we observe in spatial and temporal clustering stability are qualitatively consistent with our experimental observations of binding avidity.

### Lipid-lipid FRET measurements confirm that sphingolipid clusters are nanoscale

To further test the spatial organization of these lipid mixtures, we turned to FRET measurements that had previously established nanoscale liquid-liquid phase separation in related lipidic systems ^25, 26^. These measurements use three probes forming two donor–acceptor pairs with complementary phase preferences: naphthopyrene (Nap) partitions into ordered phases, whereas TFPC and DiD both prefer disordered phases. FRET is measured for the Nap→DiD and TFPC→DiD pairs (FRET_ND_ and FRET_TD_, respectively, Fig. S2), and we use the ratio of the two signals (FRET_R_ ≡ FRET_TD_ / FRET_ND_, Fig. 4) as the primary readout because it largely cancels temperature-dependent effects unrelated to miscibility, such as dye photophysics and changes in lipid packing (see Methods). The assay detects the onset of Lo phase upon cooling the bilayer from a uniformly mixed Ld phase; in canonical Ld + Lo mixtures, this onset appears as a breakpoint in the temperature dependence. We measured FRET in 100-nm liposomes for a positive-control 40% DPPC : 40% DOPC : 20% Chol mixture, a negative-control POPC sample, and four lipid mixtures used in our binding experiments. Relative to the same composition in micron-sized vesicles ^25^, the response in the positive control was muted: in 100 nm liposomes, domain dimensions are constrained to a few multiples of the Förster distance, so probe segregation is incomplete and the full signal change cannot develop. A breakpoint was nonetheless resolved at ∼ 35 °C, consistent with reported miscibility transition temperatures for this composition ^25, 27^. Notably, none of the tested mixtures across 0–20 mol % SM showed such a breakpoint. The 20% SM : 20% chol sample was nevertheless distinct from the others: FRET_R_ was flat or decreased slightly on cooling in the remaining mixtures, but increased steadily in this one, in the same direction as the positive control though without a resolvable transition. We interpret this as non-ideal mixing with clustering on a larger length scale than in the other mixtures, but without a true first-order transition. For the remaining mixtures, the absence of any FRET signature does not exclude heterogeneity: clustering below the ∼5-10 nm resolution of this assay would not be detected. Both readings are consistent with the simulations, in which clusters are present across the series but are largest and longest-lived at 20 mol % SM. The key finding is that these subtle variations in cluster size and persistence time are sufficient to alter viral binding by a factor of two.

**Figure 4.**
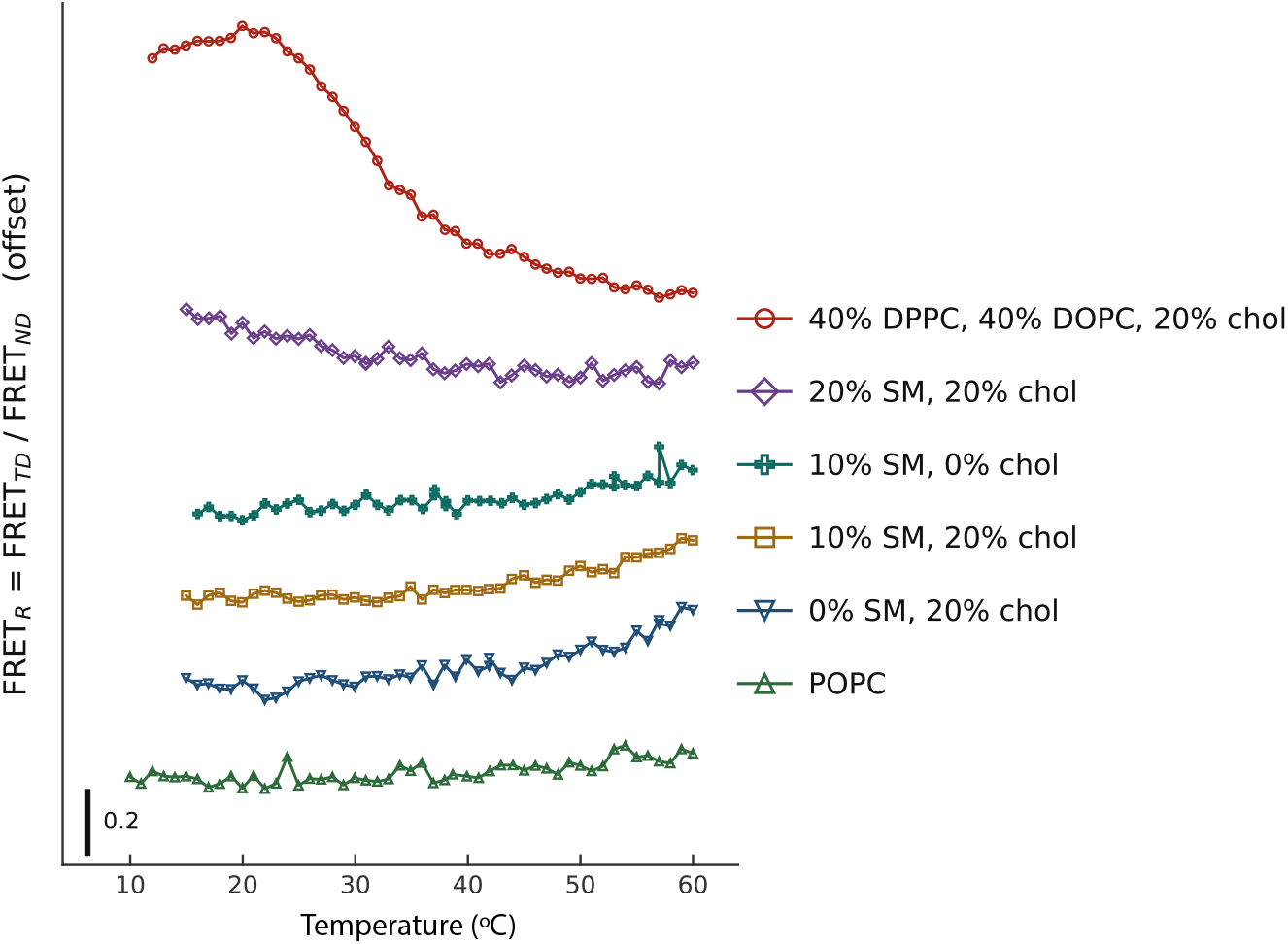
FRET measurements of miscibility in 100-nm liposomes. Steady-state FRET was measured between two donor–acceptor pairs with complementary phase preferences, Nap→DiD and TFPC→DiD, as liposomes were cooled from 60 to 10 °C in 1 °C steps (see Methods). Plotted is the ratio FRET_R_ ≡ FRET_TD_ / FRET_ND_, which largely cancels temperature-dependent effects unrelated to miscibility. The onset of Lo phase appears as a breakpoint in the temperature dependence, seen here at ∼35 °C in the 40% DPPC : 40% DOPC : 20% chol positive control but in none of the other mixtures. Of these, only 20% SM : 20% chol increases on cooling, in the same direction as the positive control. Curves are offset vertically for clarity; the scale bar denotes 0.2 in FRET_R_.

### Nanoscale clustering likely creates the environment for viral fusion

The profound enhancement of viral binding at 10 mol %, but not 20 mol %, sphingomyelin and inferred effect of nanoscale lipid clustering on viral binding led us to ask whether the resulting local inhomogeneity could persist and affect viral membrane fusion. We therefore bound influenza virus to target liposomes of different composition and then triggered fusion via a pH drop as previously described ^28^, monitoring both lipid mixing and fusion pore formation via fluorescent reporters (Fig. 5a-b). None of the compositional conditions altered lipid mixing rates or efficiencies (Fig. 5c-d), except for 5-cholesten-3-one, which had lower efficiency and slightly slower lipid mixing kinetics that could potentially be explained by the formation of gel domains in this mixture ^22^. The most interesting effects, however, were on content mixing kinetics: we observe a slowing of content mixing at 10 mol % sphingomyelin but not at 0 or 20 mol % sphingomyelin (Fig. 5e). This slowing in content mixing kinetics was both sterol- and sphingomyelin headgroup-dependent, as omitting cholesterol or replacing sphingomyelin with ceramide did not produce slowed fusion kinetics (Fig. 5f). This strongly suggests that the nanoscale lipid heterogeneity affecting viral binding persists, whether stabilized by receptor engagement or not, and alters the lipidic environment for fusion. 5-cholesten-3-one also slowed content mixing kinetics (Fig. 5f). Since liposomes containing this sterol also had altered lipid mixing (Fig. 5c), we assign this to a different mechanism than the 10 % SM : 20 % Chol mixture, but we cannot rule out additional 5-cholesten-3-one effects on fusion pore opening or the local lipid composition around fusion pores. For content mixing, omitting cholesterol greatly reduced full fusion efficiency (Fig. 5g), consistent with prior reports ^29, 30^.

**Figure 5.**
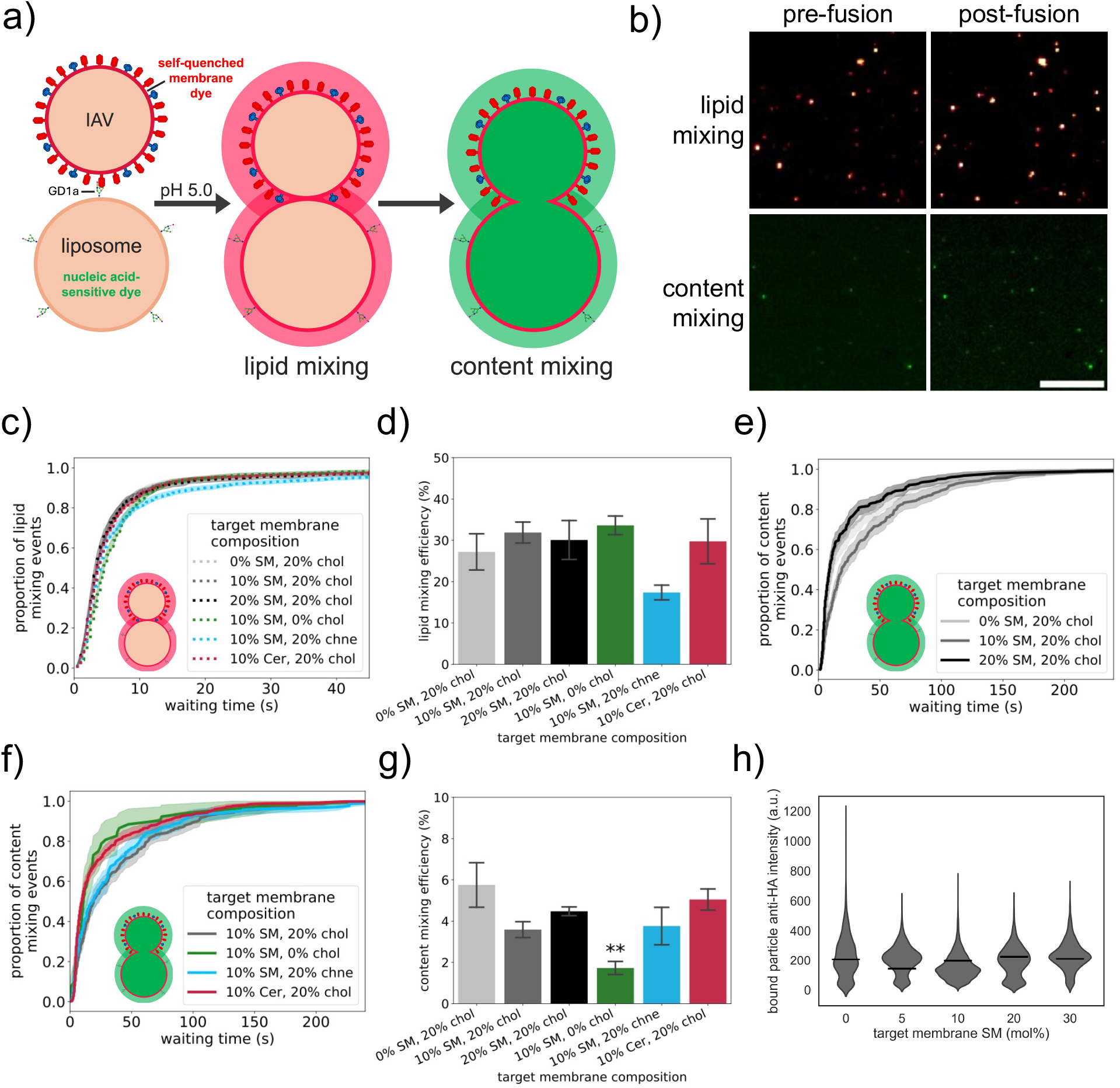
Influenza fusion with liposomes containing sphingomyelin. **a**) Schematic overview of the viral fusion assay. Fusion is triggered by buffer exchange to pH 5.0 within the flow cell channel. Lipid mixing is detected via dilutional dequenching of a lipophilic fluorophore in the viral envelope, and content mixing is simultaneously detected using a nucleic acid–sensitive dye encapsulated in target liposomes, which fluoresces after diffusing through a fusion pore and binding viral RNA. **b**) Fluorescence micrographs showing lipid (top row) and content (bottom row) mixing fluorescence before and after fusion. Each spot represents an individual fusion event. Scale bar = 10 µm. **c**) Lipid mixing kinetics of IAV with liposomes of defined lipid compositions, shown as a normalized cumulative distribution function (CDF) of lipid mixing waiting times. Only the first 45 seconds are shown for clarity. Shaded areas represent 90% bootstrapped confidence intervals. **d**) Lipid mixing efficiency, calculated as the percentage of lipid mixing events relative to the total number of fluorescently labelled IAV particles within the same field of view. **e-f**) Influenza content mixing kinetics with liposomes of differing lipid composition, plotted as normalized CDFs. Shaded areas represent 90% bootstrapped confidence intervals. **g**) Corresponding content mixing efficiencies, plotted as the percent of fluorescently labeled virions with a fluorescence increase. Statistical significance (** p < 0.01) was determined by a one-way ANOVA followed by Tukey’s HSD post hoc test. **h**) Violin plot showing anti-HA intensity of IAV particles bound to liposomes containing varying amounts of SM. Horizontal lines indicate the median anti-HA intensity. No significant differences in anti-HA intensity were observed between groups (p = 0.58 shown by a Kruskal–Wallis test). All experiments were performed at 37 °C.

If 10 mol % sphingomyelin enhances influenza binding to GD1a, it could in principle also capture virions with lower HA surface density, which may in turn fuse slower. To test whether the observed slowing in content mixing kinetics arises from such a biased subpopulation of bound viral particles, we quantified anti-HA fluorescence intensity for individual virions bound to target membranes with a range of sphingomyelin concentrations. Anti-HA intensity was independent of target membrane sphingomyelin content (Fig. 5h), indicating that 10 mol % sphingomyelin does not selectively enrich for virions with lower HA density. Instead, the observed effects are more likely attributable to changes in the physicochemical properties of the target membrane.

To confirm that the slowing in content-mixing kinetics is related to nanoscale heterogeneity in the bound membrane patch, we tested fusion to the same lipid mixtures where the GD1a receptor was replaced with a DNA-lipid tether (Fig. 6a) that localizes to Ld phases ^31^. In this binding geometry, no difference was observed in content mixing kinetics or efficiencies between IAV and 0 % SM : 20 % Chol or 10 % SM : 20 % Chol liposomes (Fig. 6b-c). This confirms that the observed slowing of fusion at 10 % SM : 20 % Chol with GD1a receptors does depend on receptor phase preference.

**Figure 6.**
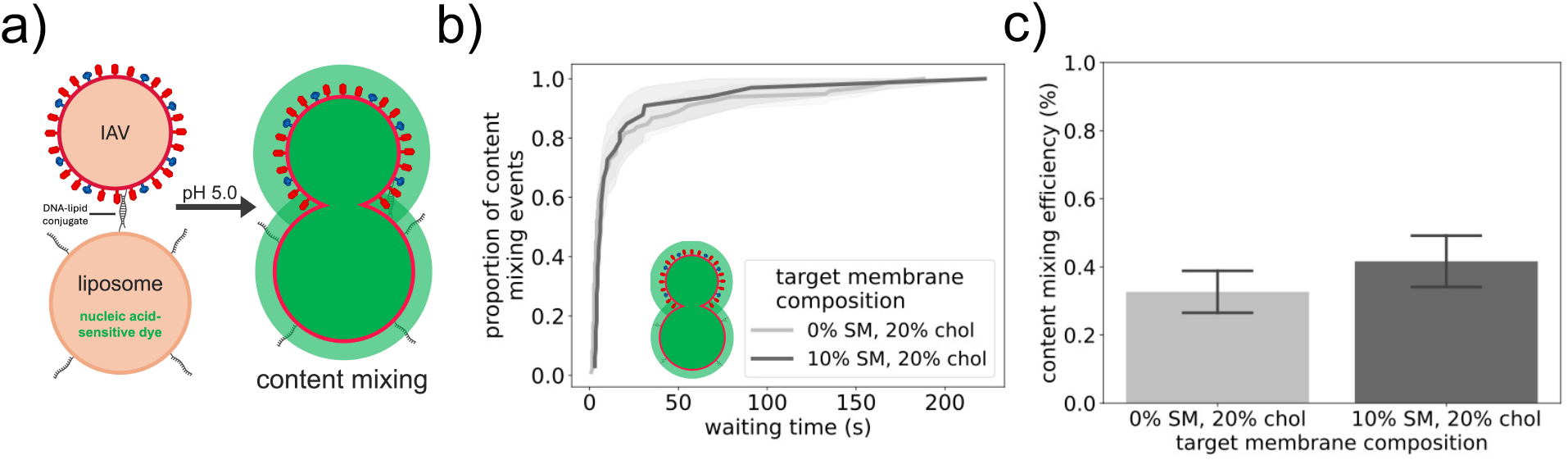
Virus bound via Ld-partitioning tethers does not show altered content mixing kinetics with sphingomyelin. **a**) Content mixing assay where viral particles are tethered to liposomes through DNA-lipid conjugates. **b**) Content mixing kinetics (similar to Fig. 5e-f) for DNA-tethered IAV. **c**) Corresponding content mixing efficiencies (similar to Fig. 5g) for DNA-tethered IAV.

### Cyclodextrin treatment reduces cell-surface binding of influenza

To test whether the role of cholesterol and sphingomyelin in promoting high-avidity influenza virus binding also applies to live cell membranes, we examined IAV attachment to A549 cells after perturbing cellular cholesterol. These studies differ from binding to model membranes in two respects: first, the plasma membrane has a much more complex and asymmetric lipid composition in addition to the full range of potential influenza receptors. Second, living cells have active trafficking and homeostatic mechanisms that may respond to perturbation. Despite the analytic complexities introduced by this second point, we believe it informative to characterize the effects of cellular cholesterol extraction on viral binding.

Although MβCD can have pleiotropic effects ^32^, it nonetheless provides a simple way to reduce plasma membrane cholesterol. We treated cells with 10 mM MßCD for 30 min at 37 ºC prior to viral binding–a relatively short treatment chosen to deplete plasma membrane cholesterol but not long enough for substantial changes to cellular lipid metabolism. Treatment with 5 mM MβCD is reported to reduce plasma membrane cholesterol by ∼50% ^33^, although the degree of depletion as well as cellular sequelae varies with dose and treatment duration. MßCD also substantially reduced cholesterol in plasma membrane vesicles in our prior work ^34^. Following lipid perturbation, IAV was bound to cells at 4 °C to prevent internalization, after which cells were fixed and immunostained for viral HA. Cholesterol depletion via MßCD significantly reduced IAV binding (Fig. 7a-b), resulting in an approximate 27% decrease in bound viral particles. Due to the multifaceted effects of cholesterol in the plasma membrane and cell trafficking responses, we cannot uniquely assign this biological effect to alterations in receptor nanoscale patterning, but it is consistent with our measurements of cholesterol effects in synthetic systems.

**Figure 7.**
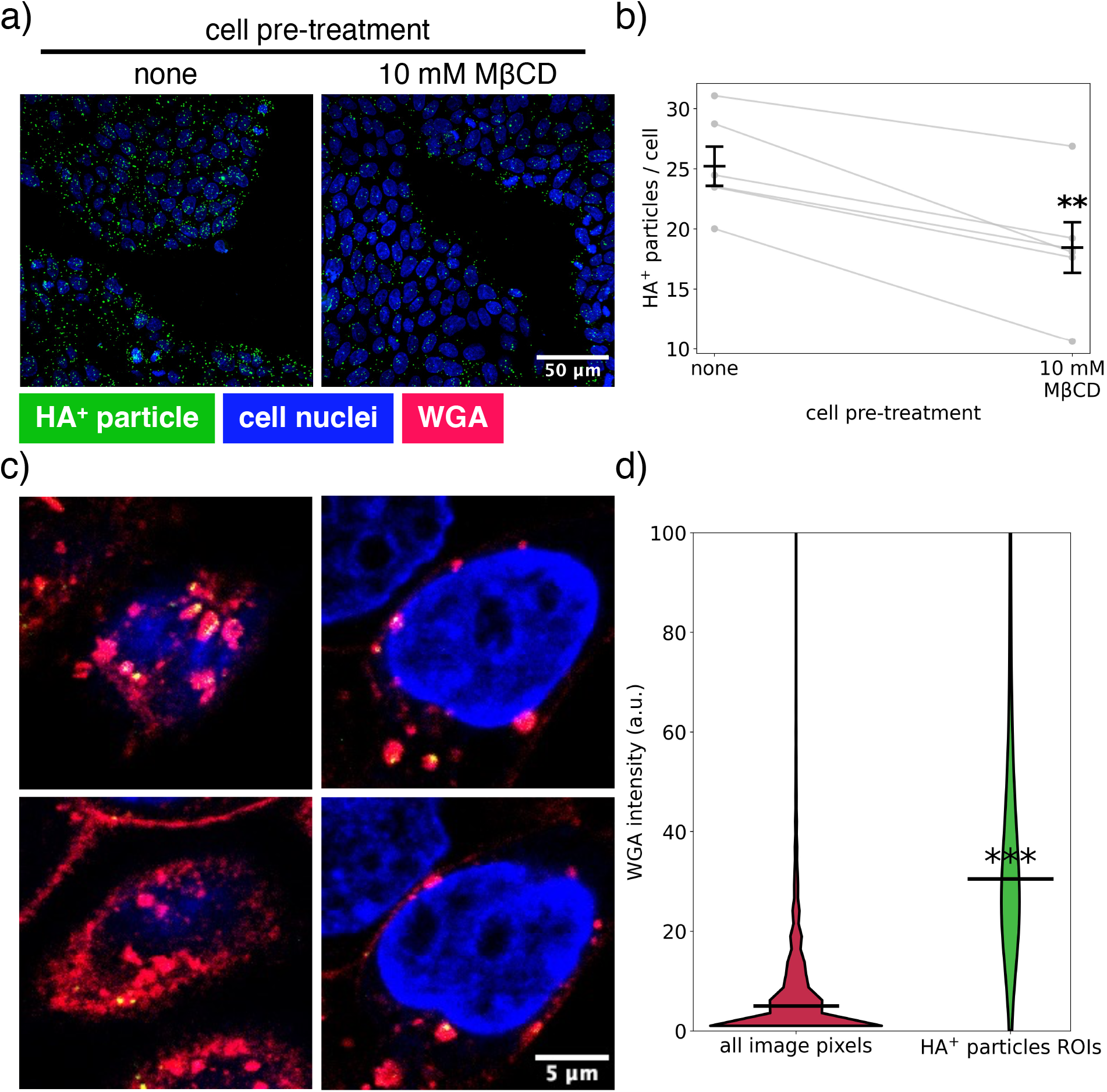
Influenza binding to cells following MβCD treatment and association of internalized influenza virus with sialylated glycans. **a**) Representative confocal micrographs of influenza virus bound to A549 cells following MβCD treatment. Images are maximum-intensity projections of confocal z-stacks. Anti-HA is indicated in green, cell nuclei in blue. **b**) Quantification of HA-positive particles per cell across biological replicates. Grey lines indicate paired measurements, and black lines show mean ± standard error of the mean. Statistical significance was assessed using a paired two-sample t-test (** p-value < 0.01). **c**) Representative confocal micrographs of internalized influenza virions (anti-HA stained, green) and sialylated glycans (WGA stained, red). **d**) Distribution of WGA pixel intensities across the entire image (red, zero-intensity pixels were excluded) and within virus-centered regions of interest (ROIs, green). Black lines indicate median intensities. The y-axis was truncated to improve visualization; only the upper tails of the violin plots are omitted. Virus-centered ROIs showed significantly higher WGA intensities than the overall image distribution (*** *p* < 0.001, Mann–Whitney *U* test).

Because we observed a relationship between sphingolipid nanoscale clustering and fusion behavior, we also wished to confirm that glycosphingolipids could persist in endosomes near the site of influenza fusion. Influenza-containing endosomes in A549 lung cell lines also stained positive with co-internalized WGA lectins (Fig. 7c-d), establishing that sialic acid ligands, and likely glycosphingolipids, were present in these endosomes. This is consistent with prior reports using fluorescently tagged GD1a that co-localizes to endosomes with other viruses ^35^. It further suggests that lipidic effects on GD1a patterning in the endocytic pathway can also affect viral fusion to endosomes.

## Conclusions

Here we show that sphingomyelin can, in the presence of cholesterol, modulate influenza binding avidity in a non-monotonic fashion. The composition yielding enhanced binding displays nanoscale heterogeneity but not macroscopic phase separation, whereas increasing sphingomyelin to levels that support macroscopic phase separation reduces binding again. Thus we hypothesize that the *spatial scale of lipid clustering controls binding avidity*. This is consistent with a model where GD1a receptors partition into Lo-like nanoclusters, and a 1:6 molar ratio of GD1a:SM at 10 mol % SM would provide a greater local enrichment of receptors than a 1:16 molar ratio in a macroscopic phase separation case at 30 mol % SM (Fig. 8). However, the precise spatial scale on which influenza virus interrogates cellular receptors remains unknown, and it is possible that these spatial scales themselves control binding rather than solely acting via receptor enrichment. The surprising finding here is that both the peak of binding avidity and the return to baseline occur below a macroscopic phase-coexistence boundary, suggesting that the effect is finely tuned. Indeed, simple modulation of the local binding equilibrium by a dilution model likely contributes but is by itself insufficient to explain the observed data. Prior work has suggested that sialic acid receptors separated by 55 Å can optimally engage adjacent binding sites on influenza virus^24^. It is thus likely that either simultaneous engagement of two sialic-acid binding sites or serial engagement at one site in a kinetic-proofreading model is responsible for the avidity peak at 10 mol % SM. The existing data do not trivially discriminate between these two possible explanations.

**Figure 8.**
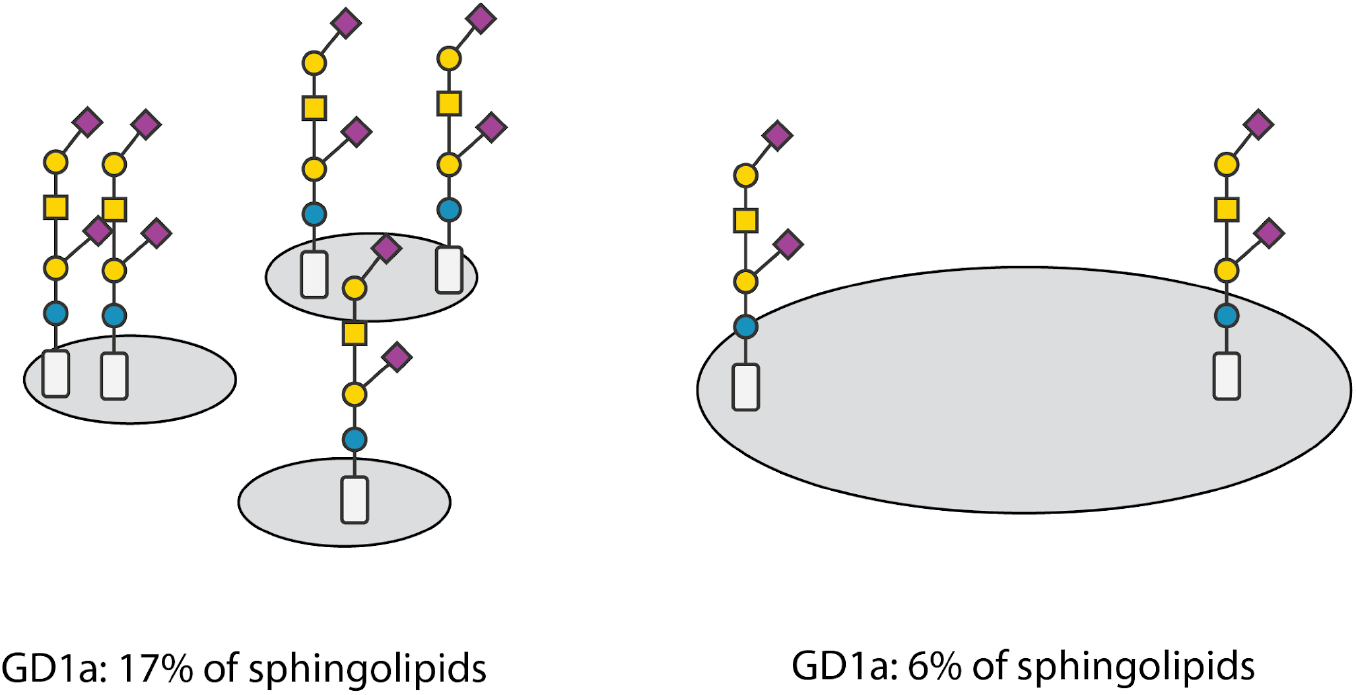
Minimal model for enhancement of influenza binding by sphingolipid domain size. Within compositions that form sphingolipid nanoclusters, GD1a approximates random assortment. As the mol % of sphingolipids is increased, GD1a is stoichiometrically diluted into a larger area. Thus small sphingolipid nanodomains will optimally increase local GD1a concentration. Mole fractions given are for 10 and 30 mol % SM.

The effects described here are in the range 5-30 mol % sphingomyelin and 20 mol % cholesterol, which is lower than the average composition of the plasma membrane outer leaflet. The observation of decreased influenza binding to cells when cholesterol is extracted suggests that cell-surface binding of influenza is nonetheless responsive to the cholesterol-sphingolipid axis. In addition, cellular activation by signaling processes can cause local plasma membrane scrambling ^36^, which would bring sphingomyelin concentrations more into the range measured here. Finally and most importantly, sphingomyelin is depleted from endosomal membranes, to the point where influenza membrane fusion may occur at a mole fraction of sphingomyelin similar to those used here. This is of particular relevance to our results suggesting that the local lipid context around GD1a receptors may influence membrane fusion -a hypothesis that is more likely if one considers both processes in the same cellular compartment. We therefore speculate that this mechanism could serve to increase influenza binding avidity within endosomes as both sphingomyelin and potentially sialic-acid receptors are depleted, maintaining viral attachment and templating the environment for viral membrane fusion.

This work uses GD1a model receptors, but a large number of other candidate influenza receptors also undergo nanoscale clustering. EGFR, a candidate receptor for influenza internalization ^7^, has been described to undergo nanoscale clustering that may be related to phase separation and can be disrupted by changing membrane cholesterol ^37^. Other glycoproteins almost certainly have non-uniform distribution in the plasma membrane ^38^, so the effects of clustering and spatial scale on viral binding avidity observed for GD1a likely generalize to other influenza receptors, although the mechanisms by which clustering are modulated may differ.

## Methods

### Viral stocks

Purified influenza A virus (strain X-31, A/Aichi/68, H3N2) was purchased from AVS Bio, and hereby referred to as IAV. All viral samples were handled under standard BSL-2 conditions with respiratory droplet precautions in accordance with approved protocols. Lipids were obtained from Avanti Polar Lipids and stored at -80 °C in chloroform.

### Liposome preparation

Lipid mixtures were prepared in glass vials at the desired mol%, followed by solvent removal under a gentle stream of nitrogen and subsequent vacuum desiccation for at least 4 h to form a dry lipid film. The dried films were rehydrated in reaction buffer (10 mM sodium phosphate, 90 mM sodium citrate, 150 mM sodium chloride, pH 7.4) pre heated at 50°C, vortexed vigorously for and incubated at 50°C for 20 minutes vortexing intermittently. For liposomes used in single-virus fusion experiments, the reaction buffer was supplemented with 100 µM of the nucleic acid-binding dye DiYO-1 (AAT Bioquest). The resulting lipid suspensions were subjected to five freeze–thaw cycles and stored at −20 °C. Prior to experiments, liposomes were thawed and extruded 21 times through 100 nm pore-size membranes. Additionally, for liposomes loaded with DiYO-1, unencapsulated fluorophore was removed using a G-25 Sephadex spin column (Cytiva). Freshly extruded liposomes were stored at 4 °C and used within one week of preparation.

### Liposome immobilization in microfluidic flow cells

Microfluidic flow cells were prepared as described previously ^28^. Liposomes were formulated to include 1 mol% 16:0 Biotinyl Cap PE to enable binding to the avidin-functionalized glass surface within the flow channel. A 10 µL aliquot of the liposome suspension was introduced into the channel and incubated overnight at 4 °C to allow surface attachment. The following day, unbound liposomes were removed by washing the channel with reaction buffer.

### IAV binding to target liposomes, viral immunolabelling and colocalization analysis

For viral binding experiments, IAV was introduced into flow channels containing immobilized liposomes with 0.5 mol% Atto 655-DOPE (ATTO-TEC) and incubated for 5 min at 37 °C to allow viral attachment. Unbound virus was removed by washing, after which the channel surface was blocked with reaction buffer containing 0.1% (w/v) bovine serum albumin. Bound virions were labelled by incubation with 1.36 µg/ml mouse anti-HA monoclonal HC3 ^39^ for 5 min, followed by washing. 2 µg/mL FITC-conjugated goat anti-mouse IgG (Invitrogen A24513) was then added and incubated for 5 min before a final wash. All washing steps were performed using reaction buffer supplemented with 0.1% (w/v) bovine serum albumin, and all antibodies were diluted in the same buffer.

Viruses bound to liposomes were imaged using 488 nm and 647 nm filter sets (Chroma) to detect FITC-labelled virions and PE-Atto 647-labelled liposomes, respectively. Images were acquired using 4% LED power, 3x3 binning, and a 2 s exposure time. Overlaid liposome and virus images were subsequently analyzed in Fiji using the ComDet plugin (https://imagej.net/plugins/spots-colocalization-comdet). The following parameters were used: maximum distance between colocalized spots = 3 pixels, approximate particle size = 1 pixel, inclusion of larger particle sizes enabled, and intensity thresholds of 2 and 10 standard deviations above background for liposomes and HA-positive particles, respectively. The percentage of liposomes colocalizing with an HA-positive particle was then quantified as a measure of IAV binding.

### Supported Lipid Bilayer Preparation and Fluorescence Recovery After Photobleaching Measurements

Glass slides were cleaned by incubation in near-boiling 7X cleaning solution (MP Biomedicals) with stirring for 30 min. Slides were then thoroughly rinsed with deionized water for 2 h and subsequently baked at 400 °C for 4 h. Polydimethylsiloxane (PDMS) squares were hole-punched and glow-discharged together with the cleaned glass slides for 30 s (15 mA, 0.39 mbar). Immediately following glow discharge, the PDMS squares were bonded to the glass slides to form wells. Liposome samples (5 µL) were added to each well within 15 min of treatment. The PDMS– glass assemblies were incubated at 37 °C for 15 min to promote liposome adsorption, rupture, and formation of supported lipid bilayers (SLBs). Excess liposomes were subsequently removed by washing the wells five times with reaction buffer. SLBs were labeled with Wheat Germ Agglutinin (WGA) conjugated to iFluor 594 (AAT Bioquest) by incubation for 30 min at room temperature, followed by an additional five washing steps. Bilayers were maintained in a hydrated state at all times and imaged within 8 h of preparation. Fluorescence Recovery After Photobleaching (FRAP) experiments were performed using a Nikon TiE microscope equipped with an iLAS2 Total Internal Reflection Fluorescence (TIRF) illuminator (Gataca systems) and a 1.49 NA 100x Apo-TIRF objective, at 37°C. Photobleaching was carried out using a 405 nm laser for 2 s over a circular spot of radius of 6.35 µm. Pre-bleach and recovery images were acquired using 561 nm excitation and a 641 nm emission filter. The TIRF angle was adjusted to generate an evanescent field depth of approximately 100 nm. For Texas Red (TR)-DHPE bleaching experiments, the same excitation and detection settings were used; however, the bleaching duration was increased to 3 s due to the higher photostability of the fluorophore.

Fluorescence recovery was analyzed using the FRAP Tools plugin in Fiji (https://imagej.net/plugins/frap-tools), followed by recovery curve generation and lipid diffusion coefficient estimation using Python code available from: https://github.com/kassonlab/FRAP_analysis_and_D_estimation. The diffusion coefficient was extracted from the FRAP data by fitting each recovery curve to the Soumpasis model ^40^. The diffusion coefficient (D) was then calculated from the fitted half-recovery time using the relation:

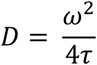

where ω is the effective radius of the bleached region and τ = half-recovery time.

### Laurdan labelling and generalized polarization (GP) measurements

Unlabeled liposomes, freshly prepared by extrusion through 100 nm pore-size membranes, were incubated with 20 µM C-Laurdan (Tocris) and shaken for 30 min at room temperature to promote dye incorporation. Laurdan-labelled liposome samples were then transferred to a black 96-well plate, and fluorescence measurements were performed at 37 °C using a Synergy H1 microplate reader. Excitation was set to 348 nm, and emission was recorded at 440 nm and 490 nm. Laurdan generalized polarization (GP) values were calculated from the emission intensities according to _41_:

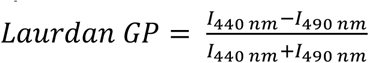

Where *I*_440 nm_ and *I*_490 nm_ represent fluorescence emission intensities at 440 nm and 490 nm, respectively. Laurdan GP provides an estimate of bulk membrane order and is sensitive to changes in global membrane packing. Higher GP values indicate increased membrane order, typically associated with reduced membrane fluidity ^41^.

### IAV lipid and content mixing assay

Prior to addition to the microfluidic flow cell channel, IAV was labelled with the lipophilic dye TR-DHPE at a self-quenching concentration as described previously ^18^. Briefly, IAV was incubated in HEPES buffer containing 15 µg/mL TR-DHPE with shaking for 2 h. Excess dye was subsequently removed by pelleting the virus and replacing the dye-containing supernatant with fresh buffer. 5µL of TR-labelled IAV was introduced to the channel containing DiYO-loaded liposomes, and incubated for 5 min at 37°C, before unbound virus was washed out. For the DNA-tethered experiments, where only content mixing was measured, 5µL of unlabeled IAV were added to the channel containing DiYO-loaded liposomes and incubated for at least 45 min at 37°C. The quantification of viral particles in the field of view was performed by viral immunolabeling as described above. The microfluidic flow cell was subsequently assembled onto the microscope stage and equilibrated to 37°C inside the environmental chamber for at least 10 min. A tubing connected to a syringe pump was attached to the outlet hole and 200µL pH 5.0 reaction buffer was added to the buffer well ^28^. Buffer exchange was initiated within the channel, and video micrographs were acquired simultaneously in the DiYO and Texas Red channels using a dual-bandpass filter set (Chroma 59022) and dichroic beamsplitter. Images were acquired for 5 min at 0.5 sec intervals, with 250 ms exposure, 4x4 binning and 4% LED power.

### IAV lipid and content mixing data analysis

Videos of fusion events were initially analyzed in Fiji using the Spot Intensity Analysis plugin (https://imagej.net/plugins/spot-intensity-analysis). Spots were detected in all frames, using a radius of 2 pixels and noise tolerance thresholds set to 100 and 30 for the lipid and content mixing channels, respectively. The resulting spot intensity–time data was subsequently analyzed in Python (code available from: https://github.com/kassonlab/single_particle_event_analysis). For each individual fluorescent particle, fusion events were identified as a sharp increase in fluorescence intensity over time. Specifically, an event was defined as an increase of at least 1000 (for lipid mixing) or 250 (for content mixing) arbitrary units, between consecutive frames, with the additional constraints that the intensity did not increase in the five preceding or decrease in the five subsequent frames by more than 25% of this value. This was to remove any transient spikes caused by fluctuations in fluorescence intensity alone. All detected lipid or content mixing events were then manually curated to remove false positive events, like when a manual focusing event occurred during image acquisition. The waiting time from fusion triggering to the observed intensity increase was calculated for each fusion event and compiled into cumulative distribution functions (CDFs). Lipid and content mixing efficiencies were calculated as the ratio of fusion events to the total number of TR-labeled IAV particles in the field of view, determined via the ComDet plugin in Fiji.

### IAV attachment and internalization assays in A549 cells

A549 cells were maintained at 37 °C in a humidified atmosphere containing 5% CO_2_ in DMEM GlutaMax (Gibco) supplemented with 10% (v/v) HyClone Cosmic Calf Serum (Cytiva) and 1% (v/v) Penicillin–Streptomycin (Gibco). Cells were seeded into clear-bottom 96-well imaging plates (Ibidi) to reach approximately 60% confluency the following day.

For the virus attachment assay, cells were treated with 10 mM MßCD diluted in Opti-MEM (Gibco) for 30 min at 37 °C. Unless otherwise stated, cells were washed thoroughly with PBS between each step. IAV, diluted in PBS, was then added, and virus attachment was carried out for 20 min at 4 °C with gentle shaking. Cells were subsequently fixed with 4% (w/v) paraformaldehyde for 15 min and stained with 1× SYTO Deep Red Nucleic Acid Stain (Invitrogen). After blocking with 0.1% (w/v) bovine serum albumin (BSA) for 25 min, bound virions were immunolabelled with 1.36 µg/mL mouse anti-HA monoclonal antibody HC3 ^39^ for 25 min, followed by incubation with 2 µg/Ml FITC-conjugated goat anti-mouse IgG (Invitrogen, A24513) for 15 min. All antibodies were diluted in 0.1% (w/v) BSA.

For the IAV internalization and sialylated glycan co-staining assay, cells were first incubated with IAV for 20 min at 4 °C as described above. Following removal of unbound virus, cells were incubated with 10 µg/mL WGA-iFluor 594 (AAT Bioquest) for 20 min at 4 °C to label cell-surface sialylated glycans. Cells were then transferred to 37 °C and incubated for 20 min in Opti-MEM containing 30 mM NH_4_Cl to allow internalization of both the virus and WGA-labelled glycans while preventing endosomal acidification and viral fusion. Cells were subsequently fixed, permeabilized with 0.1% (v/v) Triton X-100 for 10 min, and immunolabelled for influenza HA as described above.

### FRET measurements of lipid miscibility

Liposomes for FRET were prepared as described above, with naphtho[2,3-a]pyrene (Nap; TCI), TFPC (Avanti Polar Lipids) and DiD (Invitrogen) added from chloroform stock at probe-to-lipid mole ratios of 1:200, 1:1500 and 1:1000, respectively. Films containing 250 nmol total lipid were dried under nitrogen, desiccated under vacuum overnight, and hydrated in 0.500 mL HBS (10 mM HEPES, 140 mM NaCl, 2.7 mM KCl, pH 7.37) at 50 °C for 1.5 h with intermittent vortexing, followed by five freeze-thaw cycles and extrusion at 45 °C through 100 nm pore-size membranes. Immediately before measurement, liposomes were diluted 20-fold into filtered HBS that had been degassed at 60 °C for 15 min, to a final lipid concentration of 25 µM. Fluorescence was measured on a HORIBA Fluorolog-3 spectrofluorometer with a Peltier-controlled, magnetically stirred cuvette holder. Five channels were recorded at each temperature (excitation/emission, nm): Nap direct (427/484), Nap→DiD FRET (427/664), TFPC direct (499/580), TFPC→DiD FRET (499/664), and DiD direct (646/664), with 2.5 nm excitation and 5 nm emission bandpass and 0.1 s integration time. Samples were cooled from 60 to 10 °C in −1 °C steps with an initial 15 min equilibration and 2.5 min at each subsequent step. FRET in each channel pair was normalized to the geometric mean of the corresponding direct donor and acceptor signals as previously described ^25^, giving FRET_ND_ (Nap→DiD) and FRET_TD_ (TFPC→DiD), and the ratio FRET_R_ ≡ FRET_TD_ / FRET_ND_.

### Molecular dynamics simulations

Molecular dynamics simulations were computed using CHARMM36 all-atom parameters for lipids ^42^ and TIP3P water in 150 mM NaCl. Initial configurations were generated using CHARMM-GUI ^43^, and simulations are run using GROMACS ^44, 45^ with a 2-fs timestep. Long-range electrostatics were calculated using Particle Mesh Ewald, temperature was maintained at 37 °C using a velocity-rescaling thermostat ^46^, and pressure was maintained at 1 bar using a semi-isotropic stochastic rescaling barostat ^47^. Each membrane composition was simulated using 2-5 replicas for 1-2 µs per replica, and the first 500 ns was omitted from analysis.

Lo-like and Ld-like states were classified using a Hidden Markov Model where per-residue feature vectors were 6-nearest-neighbor residue identities ^48 49^. Residence times were calculated using the inferred self-transition probabilities for the Lo-like state in the model. Cluster size distributions were calculated by finding the 6-nearest-neighbors graph for all phospholipid and sphingolipid residues (using phosphorus atoms and the ceramide nitrogen as reference points) and then computing the connected subgraph sizes for sphingolipid residues. Sizes were weighted by the number of participating residues to yield a cluster probability distribution and averaged across time points and trajectories. A null distribution was estimated at each time point by randomly permuting the residue labels 10 times and measuring the permuted subgraph size distribution; values are reported as excess probability over the null distribution. Code for these analyses can be found at https://github.com/kassonlab/lipid-cluster-size-analysis.

## Acknowledgement

This work was supported by grants from the Knut and Alice Wallenberg Foundation (KAW 2020.0209) and the Swedish Research Council (2023-04674) to P.M.K. Computational studies were supported by NIH R01GM138444. Computational resources were provided by the National Academic Infrastructure for Supercomputing in Sweden (NAISS). S.M. was supported by a fellowship from the Göransson-Sandviken Foundation. The authors thank Gonzalo Sanchez for access to and assistance with FRAP imaging, the Uppsala BioVis facility for confocal microscopy, and Robert Rawle for helpful discussions.

