## Supplementary Figures for "Tuning nanoscale lipid clustering controls influenza viral binding"

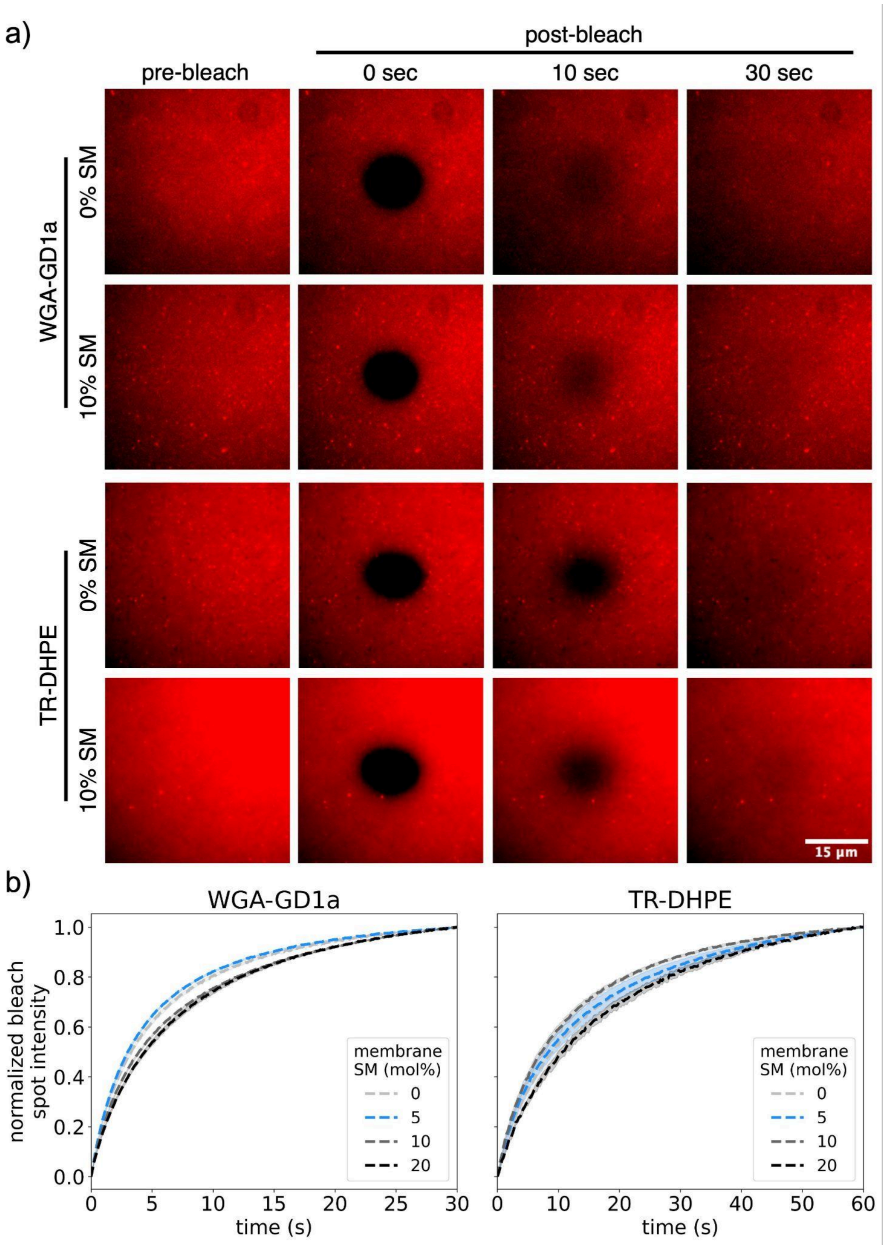


**Figure S1. Fluorescence Recovery After Photobleaching (FRAP) on supported lipid bilayers. a**) GD1a was labelled with WGA-iFluor 594 or liposomes containing 0.5 mol% TR-DHPE. A 12.7 µm diameter region was photobleached using a 405 nm laser and fluorescence recovery was subsequently monitored. Note that the bright spots correspond to non-ruptured liposomes that remained tethered to the bilayer surface. The darker spot in the upper right corner is a microscope artifact and does not affect the FRAP analysis. **b**) FRAP curves for WGA-bound GD1a or TR-DHPE, in membranes containing 0–20 mol% SM. Data were double-normalized to account for image photobleaching and pre-bleach fluorescence intensity. For both probes, fluorescence recovered to >90% of the pre-bleach intensity across all SM concentrations tested. Shaded areas represent the standard error of the mean. ≥3 independent bilayers were analyzed for each condition. Note that WGA-GD1a and TR-DHPE fluorescence recovery were imaged for 30 and 60 sec, respectively. The pre-bleach images of both WGA- and TR-DHPE-stained bilayers do not show evidence of macroscopic phase separation.

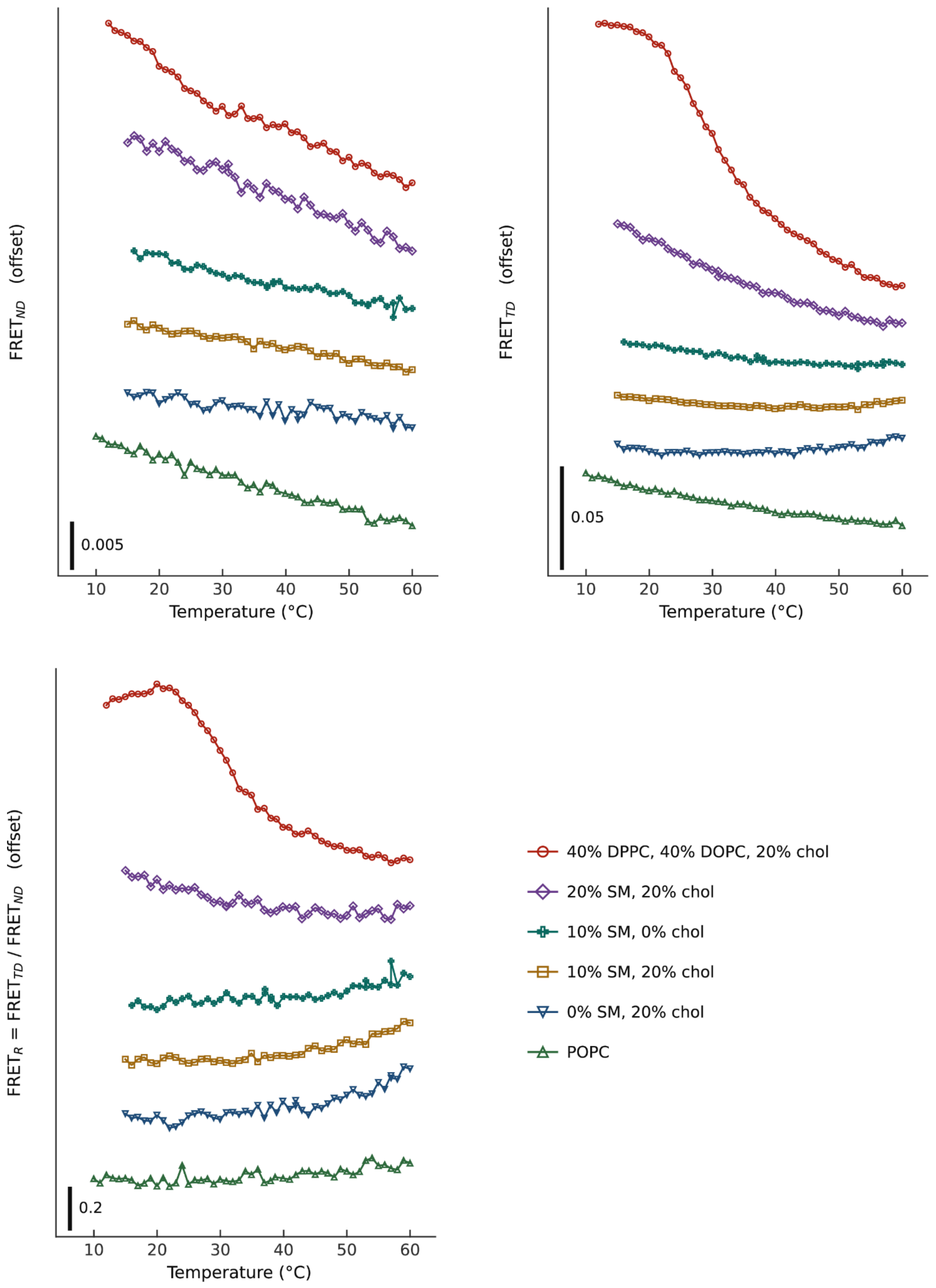


**Figure S2. Component FRET signals for lipid nanocluster analysis**. The two normalized FRET quantities Nap→DiD and TFPC→DiD pairs ($FRET_{ND}$ and $FRET_{TD}$) are plotted as a function of temperature along with their ratio $FRET_R$. The latter corresponds to the data plotted in Fig. 4.
